# Splashed E-box and AP-1 motifs cooperatively drive regeneration-response and shape regeneration abilities

**DOI:** 10.1101/2022.05.06.490982

**Authors:** Teruhisa Tamaki, Takafumi Yoshida, Eri Shibata, Hidenori Nishihara, Haruki Ochi, Atsushi Kawakami

## Abstract

Injury triggers genetic program to induce gene expression for regeneration. Several studies have recently reported the identification of regeneration-response enhancers (RREs) in zebrafish; however, it remains unclear whether a common mechanism operates in RREs. Here, we show that E-box and activator protein 1 (AP-1) motifs cooperatively function as RREs. We identified three RREs from the *fn1b* promoter by a search of conserved sequences and an in vivo transgenic assay for regeneration-response in zebrafish. Two of them derived from transposons displayed RRE activity only when combined with the −0.7 kb *fn1b* promoter, while another non-transposable element functioned as a standalone enhancer. A search for transcription factor-binding motifs and validation by transgenic assay revealed that both of E-box and AP-1 motifs are necessary and sufficient for RREs. Such RREs responded to variety of tissue injuries including zebrafish heart and *Xenopus* limb bud regenerations. Our findings highlight that regeneration is regulated by merging two activating signals evoked by tissue injuries. It is speculated that a large pool of potential enhancers in the genome shaped regenerative capacities during evolution.

**SUMMARY STATEMENT:** The study revealed that regeneration-response enhancer is composed of two transcription factor-binding motifs. The fidelity of regeneration-dependent gene expression is ensured by merging two activating signals evoked by injuries.

## INTRODUCTION

Multicellular organisms maintain tissue integrity by regenerating or repairing injured parts. However, this ability differs depending on the species, tissue, and developmental stage (Kumar et al., 2007; Kawakami, 2010; Poss, 2010; Tanaka, 2016). Such heterogeneity could be due to the loss of key genes during evolution (Ivanova et al., 2013); however, changes in expression of key genes are likely to be the main cause of differing regenerative capacity because this ability is not specific to certain evolutionary clades (Bely and Nyberg, 2010).

During regeneration, cells sense injuries and initiate the expression of related genes, where the distal-acting regulatory sequences, enhancers, play a pivotal role (Visel et al., 2009). However, the identification of enhancers for complex biological processes, such as regeneration, has been hampered by the need for *in vivo* assays. Recently, several regeneration-response enhancers (RREs) have been reported in zebrafish (Kang et al., 2016; Pfefferli and Jaźwińska, 2017; Wang et al., 2020; Goldman and Poss, 2020), and it has also been suggested that such enhancers and their response to injury are conserved in hearts and fingertips of neonatal mice (Kang et al., 2016). Interestingly, studies so far suggested that the activator protein 1 (AP-1)-binding motifs are necessary for RRE activity. However, AP-1, a heterodimeric protein consisting of Jun and Fos, is known to be essential for a variety of biological processes (Hess et al., 2004). Therefore, it remains unclear whether the AP-1 signal is sufficient for initiating the precisely regulated regeneration process.

Here, we identified RREs through the search of conserved sequences around the regeneration-induced genes and the in vivo transgenic assay for amputation-response, while many of recent studies have adopted high-throughput approaches of chromatin accessibility analyses for searching RREs (Kang et al., 2016; Lee et al., 2020; Thompson et al., 2020; Wang et al., 2020). From comparison of RREs, we found that the regeneration response enhancer, RRE, is composed of two TF-binding motifs, E-box and AP-1. Our study highlighted that the fidelity of regeneration-induced gene transcription is ensured by merging two different signals activated by injuries.

## RESULTS AND DISCUSSION

### Conserved sequences around regeneration-response genes

During the regeneration process, signals activated by injuries reach the respective enhancers to initiate gene expression. To elucidate the enhancer elements, we searched for conserved sequences between the bacterial artificial chromosome (BAC) sequences of fn1b (Yoshinari et al., 2009; Shibata et al., 2018) and msxc (Akimenko et al., 1995), which drive regeneration-dependent EGFP expression in the wound epidermis (WE) and blastema, respectively. We found highly conserved sequences, termed as E1 and E4, which showed > 60 % nucleotide identity between the respective homologous sequences. Additional conserved sequences, E5 and E6, were also discovered by searching the BAC clones of other regeneration-induced genes, *junba, junbb*, and *dlx5a* (Yoshinari et al., 2009). Multiple copies of conserved sequences were distributed within and around the regeneration-response genes (Fig. S1A).

### Conserved sequences are derived from transposable elements

Considering the high sequence homology in non-coding region, we suspected that the conserved sequences were derived from transposable elements (TEs). Indeed, the repeat annotation in the UCSC Genome Browser indicated that all conserved sequences were derived from TEs such as short interspersed elements (SINEs) and non-autonomous DNA transposons (MITEs) (Fig. 1B). E1, E4, E5, and E6 sequences diverged 18–26 % (including indels) or 12–15 % (excluding indels) of sequences from each TE consensus. RepeatMasker analysis of the zebrafish genome showed that the number of each TE copy in the genome was 16,298 (E2), 43,792 (E4), 11,391 (E5), and 13,789 (E6), many of which diverged by > 10 % (Fig.1B; Fig. S1B, C). Since it is estimated that a 7 % divergence of human TEs corresponds to an insertion around 50 million years ago (Lander et al., 2001), many of these TEs were thought to be inserted in the genome more than ten million years ago.

**Fig. 1.**
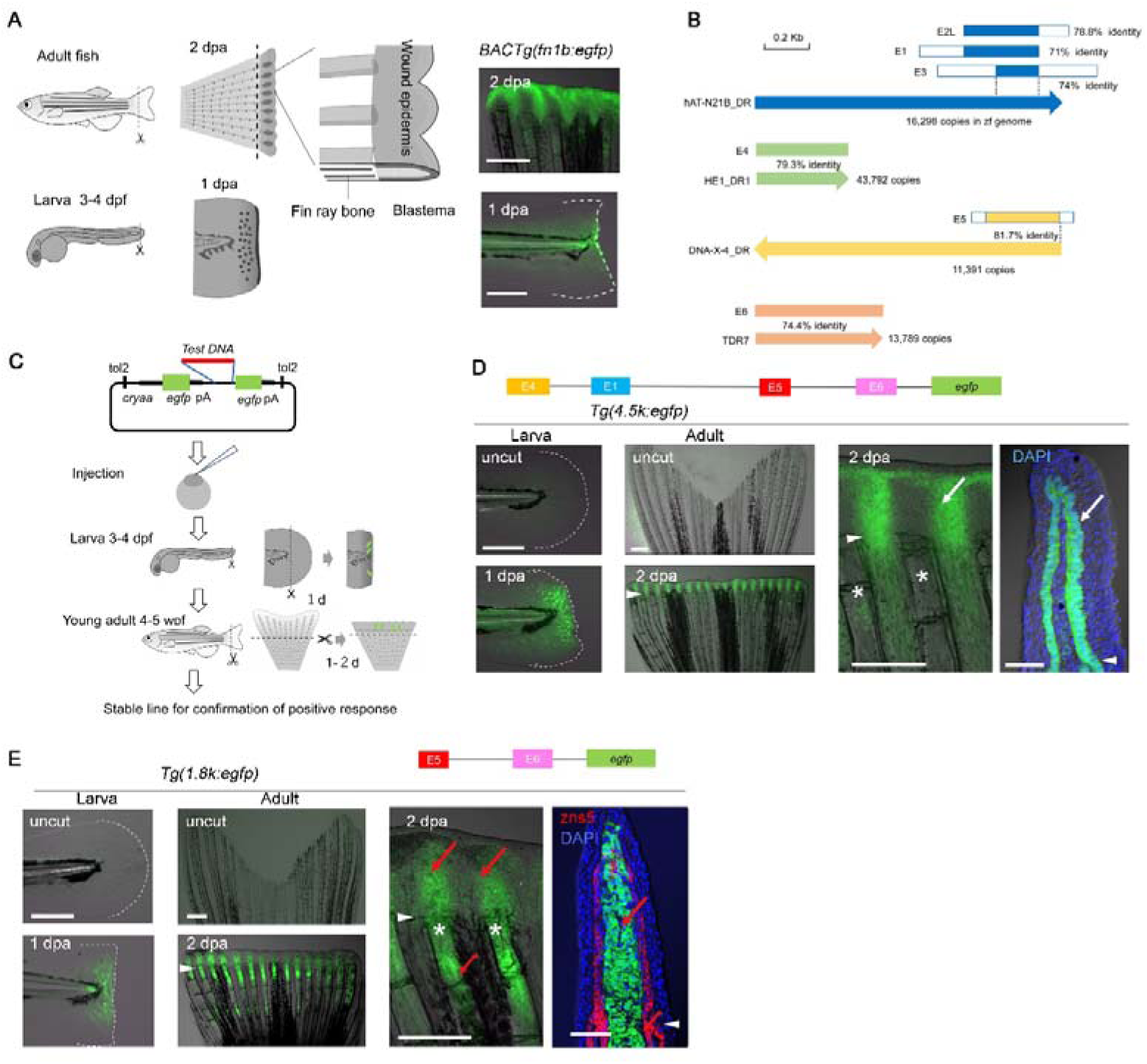
Regeneration-dependent response of the *fn1b* Promoter. (A) Regeneration of adult fish fin and larval fin fold, and amputation-induced EGFP expression in the *BAC Tg(fn1b:egfp)*. (B) Conserved DNA elements in the *fn1b* promoter that have homology to TEs. hAT-N21B_DR, DNA-X-4_DR and TDR7 are types of MITEs. HE1_DR1 a type of SINEs. These TEs exist in the zebrafish genome more than 10,000 copies. (C)Transgenic EGFP reporter assay procedure for testing RRE activity. (D, E) Amputation-dependent EGFP expression in the larval fin fold (left panels) and adult fin (middle panels) of *Tg(4*.*5k:egfp)* (D) and *Tg(1*.*8k:egfp)* (E). Right panels show higher magnification of the adult fin. zns5, antibody staining against regenerating osteoblasts. EGFP expression was confirmed in two Tg lines, respectively. Arrowhead, amputation plane. Asterisk, fin ray. White arrow, EGFP in the basal epidermis; red arrow, EGFP in the fin ray mesenchyme and blastema. Scale bars: 50 µm (left panels), 500 µm (middle panels), and 100 µm (sections).

### RRE activity contained in the *fn1b* promoter

In mammals, it has been shown that several TEs possess cis-regulatory functions and serve as tissue-specific enhancers (Chuong et al., 2016; Nishihara, 2019). To test whether these conserved sequences have an RRE activity, we performed an EGFP reporter assay. Constructs with test sequences placed upstream of the *egfp* cassette were injected into fertilized eggs and assayed for their RRE activity during regeneration in the larval fin fold (Kawakami et al., 2004) and in the young adult fin of the F0 animals (Fig. 1C). For constructs that showed a positive response in the F0 assay, the activity was validated in more than two Tg lines to avoid a possible positional effect of the integrated genomic site.

Firstly, we tested the *Tg(4*.*5k:egfp*), which contains the E1, E4, E5, and E6 Tes upstream of *fn1b*, and found that it induced EGFP in response to amputation of the fin fold and fin (Fig. 1D; Fig. S2A). Regeneration-dependent EGFP expression was localized mostly in the basal layer of WE and also in a small number of fin ray mesenchymal cells proximal to the amputation plane as in *BAC Tg*(*fn1b:egfp*) (Fig. S2B). The response was also observed in the −3.2 kb and −1.8 kb Tgs (Fig. 1E and S2C–E), but further truncated regions, −0.8 kb and −0.4 kb, did not display RRE activity (Fig. S2C), suggesting that an RRE exists at least between −1.8 kb and −0.8 kb that contains E5. Interestingly, while the −4.5 kb and −3.2 kb promoters drove the EGFP localization in the WE, the expression in the −1.8 kb promoter Tg shifted to the fin ray mesenchymal cells that give rise to the blastema (Fig. 1E), implying the presence of an element that promotes WE-specific expression between −3.2 kb and −1.8 kb.

### Identification of RREs within TEs and in a non-TE region

To examine whether E5 contains an RRE, the construct *E5-0*.*7k*, in which E5 was directly placed upstream of the −0.7 kb promoter, was tested. This construct displayed RRE activity in the mesenchymal cells like the −1.8 kb Tg (Fig. 2A; Fig. S2F, G), indicating that E5 indeed contains an RRE. However, this response was not detected in the construct where E5 was placed upstream of the short *minimum promoter* (*miniP*) (Fig. S2F). Therefore, it is suggested that E5 functions as an RRE only if it is placed in combination with a helper element within the −0.7 kb promoter. We further tested the RRE activity of E4 and found that E4 also contains an RRE that requires a helper element within the −0.7 kb promoter (Fig. 2B; Fig. S2F, H).

**Fig. 2.**
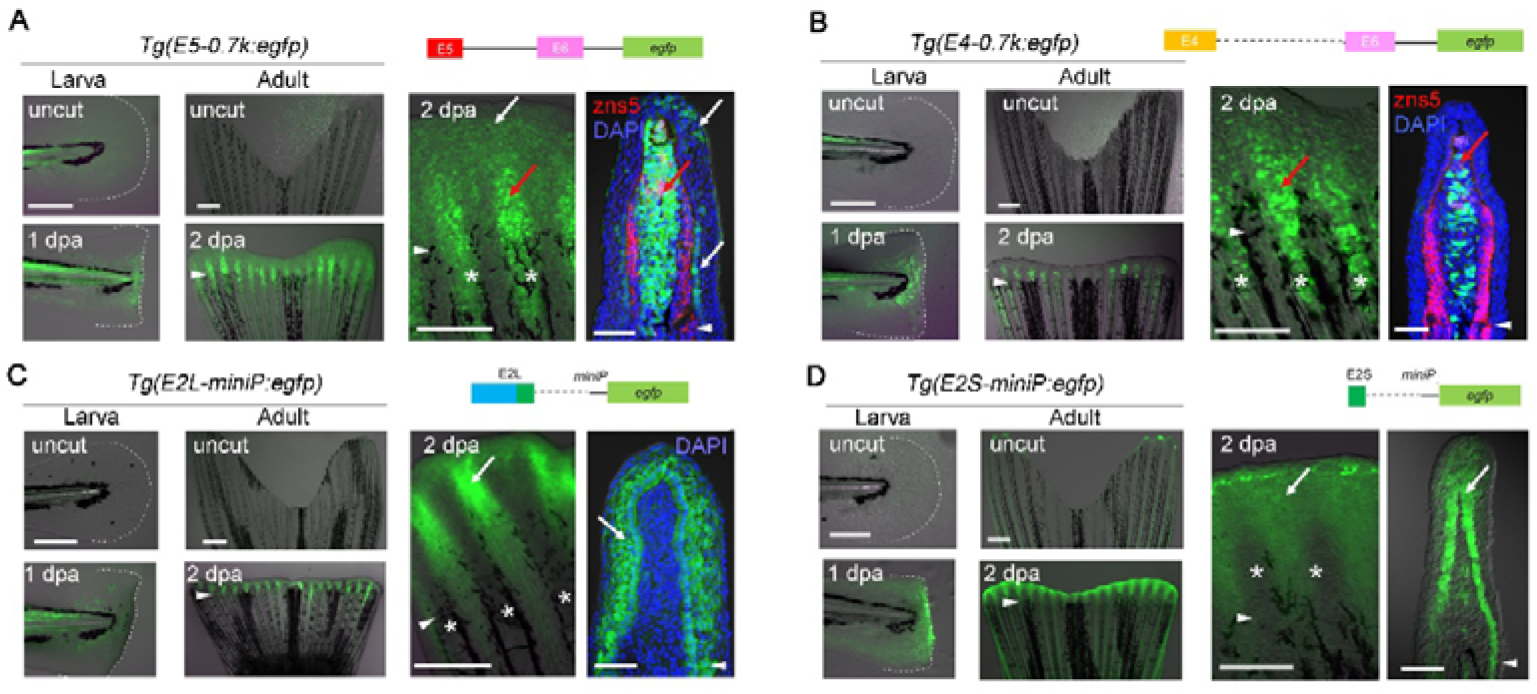
RRE activities in TEs and non-TE Sequences. (A-D) Regeneration-dependent EGFP expression in the larval fin fold (left panels) and adult fin (middle panels) of the transgenic lines, *Tg(E5-0*.*7k:egfp)* (A), *Tg(E4-0*.*7k:egfp)* (B), *Tg(E2L-miniP:egfp)* (C), and *Tg(E2S-miniP:egfp)* (D). Right panels show higher magnification of the adult fin. zns5, regenerating osteoblasts. EGFP expression was confirmed in two Tg lines, respectively. Note that E2S does not contain the E2 TE sequence. Arrowhead, amputation plane. Asterisk, fin ray. White arrow, EGFP in the basal epidermis. Red arrow, EGFP in the fin-ray mesenchyme and blastema. Scale bars: 50 µm (left panels), 500 µm (middle panels), and 100 µm (sections).

To further investigate the presence of RREs in other TEs, we constructed E2L, which contains E2 TE and a short flanking 100-bp genomic sequence. Interestingly, it displayed RRE activity in combination with *miniP* and drove EGFP expression in WE (Fig. 2C; Fig. S2F, I), indicating that E2L functions as a standalone RRE. More significantly, when we tested E2 itself, E2 did not display RRE activity, but the short flanking non-TE sequence (E2S, ∼100 bp) functioned as an RRE (Fig. 2D; Fig. S2F, J). Together, a series of Tg assay revealed that the upstream *fn1b* promoter region contains at least 3 RREs.

### E-box and AP-1 motifs are contained in RREs

Thus, we have identified that E4, E5 and E2S contain RREs, and that the *fn1b* −0.7kb region, which includes E6, has a helper element that facilitates the response of E4 and E5. To identify the responsible transcription factor (TF)-binding sites, we searched for the motifs that were shared between these elements.

On comparing E2S, E4, and E5, we realized that highly conserved E-box motifs were commonly contained (Fig. 3A, C; Fig. S3A). The motif has consensus core sequences, CAGCTG or CACCTG, and is the binding target of basic helix-loop-helix (bHLH) TF factors such as Tcf, MyoD, Mesp, Twist, Clock, and Myc (Jones, 2004; Wang and Baker, 2015). In E4, the clustered E-box motifs were located within the conserved 3′-tail domain of the TE (Fig. S3B), suggesting that E-box motifs may be preserved in a significant population of HE1_DR1.

**Fig. 3.**
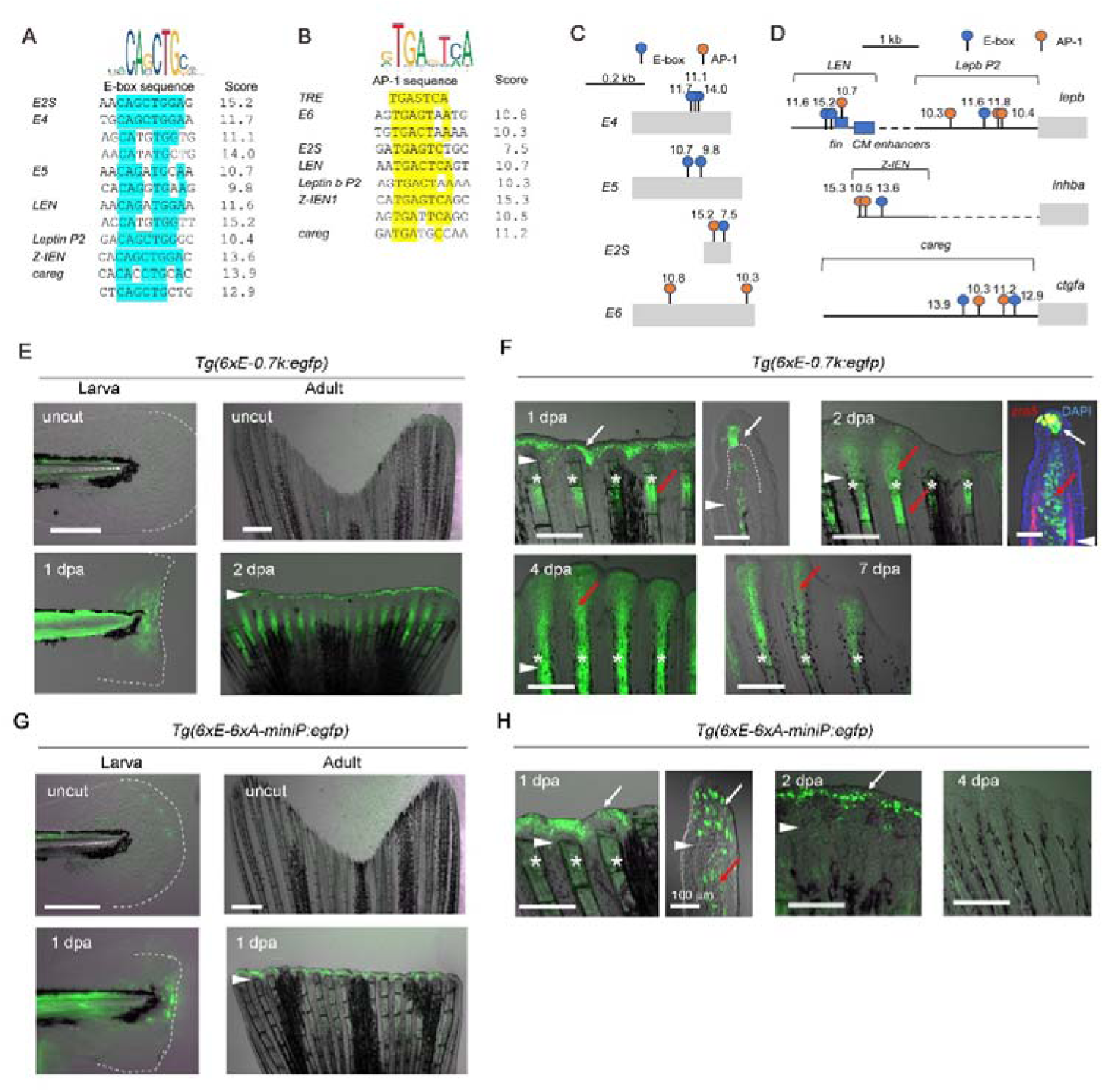
Combination of E-box and AP-1 motifs acts as the RRE. (A, B) Alignment of E-box (A) and AP-1 (B) motifs found in the RREs of *fn1b, lepb* (*LEN*), *inhba* (*IEN*), and *ctgfa* (*careg*). TRE, a consensus 12-O-tetradecanoylphorbol-13-acetate responsive element. Only the respective motifs with higher Jasper scores (> 10.0) are shown, except for those of the E-box in E2S and E5, which are slightly lower. (C, D) Distribution of the E-box and AP-1 motifs in various RREs. Numbers above the E-box and AP-1, the Jasper scores. (E) Regeneration-dependent EGFP expression in the larval fin fold (left panels) and adult fin (right panels) of the transgenic line *Tg(6*×*E-0*.*7k:egfp)* (n = 3 Tg lines). (F) EGFP expression in *Tg(6*×*E-0*.*7k:egfp)* at different regeneration stages of the adult fin. The right panels of 1 and 2 dpa are sections. zns5, regenerating osteoblasts. (G) Regeneration-dependent EGFP expression in the larval fin fold (left panels) and adult fin (right panels) of the transgenic line *Tg(6*×*E-6*×*A:egfp)* (n = 2 Tg lines). (H) Regeneration-dependent EGFP expression in *Tg(6*×*E-6*×*A:egfp)* at different regeneration stages of the adult fin. Arrowhead, amputation plane. Asterisk, fin ray. White arrow, EGFP in the basal epidermis. Red arrow, EGFP in the fin ray mesenchyme and blastema. Scale bars: 50 µm (larvae), 500 µm (adult fin), and 100 µm (adult fin section).

Furthermore, the comparison of TF-binding sites between E2S and *fn1b* −0.7kb showed that AP-1 motifs are common between E2S and E6 (Fig. 3B, C; Fig. S3A). AP-1 is a group of leucine zipper TFs that are composed of heterodimers of Fos, Jun, and other families of proteins, and binds to the consensus core sequence TGASTCA (Hess et al., 2004). Significantly, highly conserved E-box and AP-1 motifs were also found within the RRE regions of the previously identified RREs, *LEN* (Kang et al., 2016), *Z-IEN* (Wang et al., 2020), and *careg* (Pfefferli and Jaźwińska, 2017) (Fig. 3D), supporting the notion that E-box and AP-1 motifs are common TF motifs in RREs.

### Combination of E-box and AP-1 motifs act as RRE

To validate whether the E-box and AP-1 motifs are functionally required for regenerative response, the respective TF motifs were removed from E2S, and the response was examined. In the E2S(dE) and E2S(dA) constructs, in which the E-box and AP-1 motifs were removed, respectively, the amputation-induced EGFP expression was abolished (Fig. S2F), demonstrating that both of E-box and AP-1 motifs are essential for regenerative response.

We further assayed the regenerative response of one of copies of HE1_DR1, which had a divergent sequence in the E-box cluster (Fig. S3B). Unlike E4, the HE1_DR1(m) construct did not display a regenerative response (Fig. S3C), indicating that not all the HE1_DR1 TEs retain the RRE activity.

To further prove that these TF motifs are sufficient for the regenerative response, we created constructs that contained tandem repeats of the E-box and/or AP-1 motif and examined the response. Although the constructs with either E-box or AP-1 did not show a response (Fig. S3D), the constructs that contain both E-box and AP-1 motifs, *6*×*E-0*.*7k* and *6*×*E-6*×*A-miniP*, displayed a response to amputation (Fig. 3E–3H; Fig. S3E, F). Furthermore, the mutated E-box construct displayed a severely decreased response (Fig. S3D). From these results, we concluded that the combination of E-box and AP-1 motifs is sufficient for a regenerative response.

Interestingly, the localization of EGFP induction differed between the constructs: EGFP expression was detected in the distal WE in *Tg(6*×*E-6*×*A-miniP:egfp*), whereas it was detected in both the distal WE and mesenchymal cells that give rise to the blastema in *Tg*(*6*×*E-0*.*7k:egfp*) (Fig. 3F and 3H). It is thought that this difference may be due to the expression of responsible bHLH and/or AP-1 factors that can bind to the TF motif sequences in the respective constructs.

Though we found three RREs in the *fn1b* promoter, it is thought that they may be functionally redundant. This is possibly why amputation-dependent gene expression could not be completely abolished by removing one or several RREs from the genome (Thompson et al., 2020).

### RRE responds to injuries in variety of tissues and other species

It has been shown that an incision in the inter-ray tissue does not induce a regenerative response, but that an injury to the fin ray induces a regenerative response on the proximal side of the injury (Gauron et al., 2013; Wang et al., 2020). To validate whether the response mediated by the E-box/AP-1 RRE was a regenerative response, we performed a similar assay using the WE-responding line *Tg(E2L:egfp)* and the mesenchyme-responding line *Tg(E4-0*.*7k:egfp)*. Both lines displayed EGFP induction by ray injury but not by inter-ray incision (Fig. 4A), confirming that the E-box/AP-1 RRE is only activated by the regenerative signal.

**Fig. 4.**
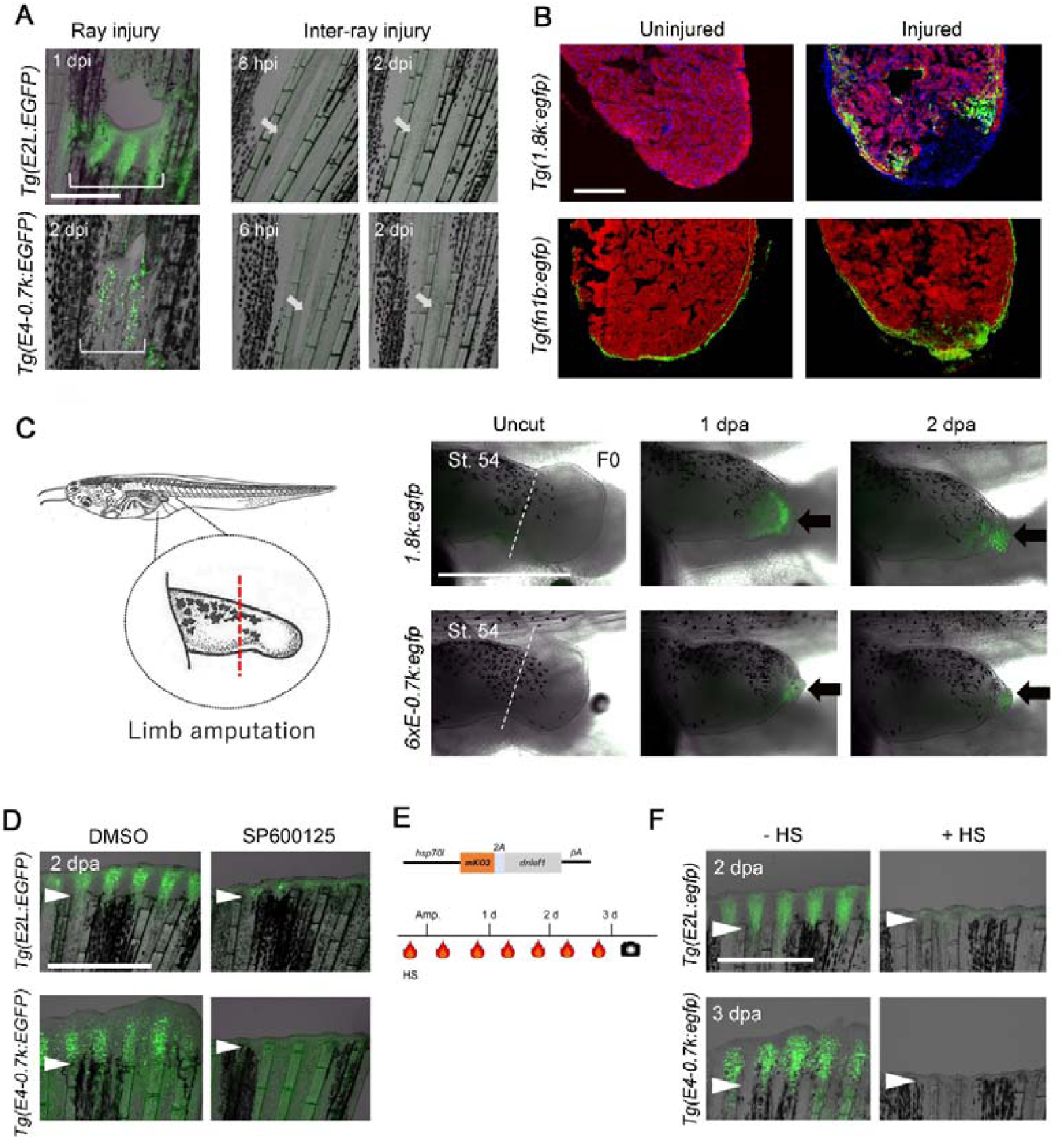
Conservation of E-box/AP-1-mediated regeneration response beyond tissue types and species. (A) EGFP expression induced by ray injury, but not by inter-ray incision in *Tg(E2L:egfp)* (upper panels) and *Tg(E4-0*.*7k:egfp)* (lower panels). EGFP induction was only detected on the proximal side of the injury. Arrow, site of inter-ray injury, which rapidly healed within 2–3 dpi. Scale bar: 500 µm. (B) Induction of EGFP expression in *Tg(1*.*8k:egfp)* (upper panels) and *BAC Tg(fn1b:egfp)* (Shibata et al., 2018) (lower panels) after heart resection. Cryosections were stained with anti-EGFP, MF20 for cardiac muscles, and DAPI. *Tg(1*.*8k:egfp)* showed EGFP induction in the cardiomyocytes around the amputation site after heart resection (n = 2 fish for uninjured and injured, respectively). The EGFP expression in the epicardial cells was seen in uncut heart of the *BAC Tg(fn1b:EGFP)* (n = 3 fish), but it was upregulated by heart resection (n = 2 fish). Dotted line, amputation site. Scale bar: 200 µm. (C) Detection of EGFP expression in the amputated tadpole limb bud of *Xenopus laevis*, which was injected with the respective constructs at the one-cell stage. Arrows, EGFP expression. Dotted line, site of amputation. EGFP response was observed in mesenchymal cells (*1*.*8k:egfp*, n = 12 of 12 with lens EGFP^+^ tadpole) and the distal tip of epithelial cells (*6xE-0*.*7k:egfp*, n = 8 of 8 with lens EGFP^+^ tadpole). Scale bar: 1 mm. (D) Knockdown of the EGFP induction by the JNK inhibitor SP600125 in *Tg(E2L:egfp)* (n =10) and *Tg(E4-0*.*7k:egfp)* (n =10), respectively. Arrowhead, amputation plane. Scale bar: 1 mm. (E) dnLef1 construct and the experimental procedure. (F) Knockdown of EGFP expression of *Tg(E2L:egfp)* (n = 3) and *Tg(E4-0*.*7k:egf)* (n = 12) by dnLef1. Arrowhead, amputation plane. Scale bar: 1 mm.

We further examined the response of the E-box/AP-1 RREs in other tissues. The pectoral paired fins displayed an EGFP response like that in the tail fin (Fig. S4A). EGFP expression was also detected during scale regeneration (Fig. S4B). Furthermore, the zebrafish larvae that were amputated at the trunk region through the neural tube, somites, and notochord, were unable to completely regenerate; however, EGFP expression was also induced in the epithelial tissue of *Tg(E2L:egfp)* and in the mesenchymal cells of *Tg(E4-0*.*7k:egfp)* (Fig. S4C).

Furthermore, we examined the response of E-box/AP-1 RRE during heart regeneration using the *Tg(1*.*8k:egfp)*. The EGFP expression was induced in the cardiomyocytes around the amputation site after heart resection, while no EGFP fluorescence was detected in the uninjured heart (Fig. 4B), indicating that the RRE also directs regenerative response in the heart. Interestingly, in contrast to the EGFP expression in the cardiomyocytes, the *BAC Tg(fn1b:egfp)* (Shibata et al., 2018) displayed EGFP expression in the epicardial cells as in the endogenous *fn1b* expression (Wang et al., 2013).

We further asked whether the function of E-box/AP-1 RRE is evolutionary conserved in other species. We introduced the constructs, *1*.*8k:egfp* or *6*×*Ebox-0*.*7k:egfp*, into *Xenopus laevis*, an animal that displays incomplete regenerative abilities depending on the developmental stage (Phipps et al., 2020), and examined the response during limb bud regeneration. In injected tadpoles, EGFP induction was detected in the amputated limbs (Fig. 4C), suggesting a conserved mechanism of E-box/AP-1 RRE beyond species.

### Both AP-1 and E-box signals are required for RRE response

We further investigated whether the signals through the E-box/AP-1 RRE are essential for the transcriptional response and regeneration. It has been suggested that Junba and Junbb, components of AP-1, are phosphorylated by the Jun N-terminal kinase (JNK) to play an essential role in fin regeneration (Ishida et al., 2010), and that the JNK inhibitor, SP600125, knockdowns EGFP induction in the Z-IEN RRE (Wang et al. 2020). We tested the effect of SP600125 on *Tg(E2L:egfp)* and *Tg(E4-0*.*7k:egfp)* and observed that amputation-induced EGFP expression is downregulated by the inhibitor (Fig. 4D).

We further knocked down the E-box-mediated signalling by expressing the dominant negative form of Lef1 (dnLef1) under control of the *heat shock promoter* (*hsp70l*) (Fig. 4E). Among the members of the bHLH protein family, a typical E-box motif, 5′-CAGCTG-3′, is bound by the E2A subgroup including Tcf3 and Tcf12 (Cadigan and Waterman, 2012). The Tcf/Lef family proteins have a β-catenin-binding domain in the N-terminus, and the truncated Lef1 acts as a dominant-negative (Vacik and Lemke, 2011). We validated the effect of dnLef1 during embryonic development and confirmed that the expression of exhibited severe developmental abnormalities that resemble to those observed in decreased β-catenin signalling (Hao et al., 2013) (Fig. S5). When dnLef1 was overexpressed in *Tg(E2L:egfp)* or *Tg(E4-0*.*7k:egfp)* during regeneration, EGFP induction was diminished, accompanied by complete suppression of regeneration (Fig. 4F). These results suggest that the Tcf/Lef signal through the E-box motif is essential for transcriptional response and regeneration.

## Conclusions

Regeneration-dependent enhancers have recently been identified in zebrafish and killifish (Kang et al., 2016; Pfefferli and Jaźwińska, 2017; Lee et al., 2020; Thompson et al., 2020; Wang et al., 2020); however, whether these enhancers function via a common mechanism has not been clarified yet. Here, we identified multiple RREs from the *fn1b* promoter region and revealed that the regeneration-response enhancer, RRE, is composed of two TF-binding motifs, E-box and AP-1, and that the signals through both motifs cooperatively activate gene expression. Our study demonstrates that E-box and AP-1 RRE mediate the regeneration-induced transcriptional response in variety of tissues and in other species. From these data, we propose that E-box/AP-1 is a universal regeneration-response machinery (Fig. S6A).

As discussed in the previous paper (Wang et al., 2020), the AP-1 complex has an ancient evolutionary origin and may have an original function in priming the injury response. The *jun* and *fos* gene families are known as early response genes that mediate an early signal following tissue trauma. We speculate that additional signal mediated by the E-box motif, which may detect a different aspect of tissue integrity, such as tissue loss, was acquired later in evolution to ensure correct and faithful regulation of the regenerative response.

Although it is still unknown which Tcf and AP-1 proteins are responsible for the regenerative response, it is hypothesized that different members of Tcf and/or AP-1 could be involved in different tissues depending on their expression in respective tissues. Tcf and AP-1 proteins and their specificity to the heterogeneous E-box and AP-1 sequences may determine the responding tissue. Interestingly, this also implies that epidermal and mesenchymal tissues independently respond to regeneration.

The function of TF motifs in the TE life cycle is still debatable (Hermant and Torres-Padilla, 2021), but potential TF motifs have been splashed throughout the genome during the divergence of species and expansion of TEs (Nishihara, 2020) to form a huge reservoir of potential enhancers. Such motifs could repeatedly gain and lose RRE activity by mutations through the course of evolution, and the resulting selection of responsive genes may have shaped an organism’ s ability to regenerate (Fig. S6B).

## MATERIALS AND METHODS

### Zebrafish maintenance

All fish were maintained in a recirculating water system in a 14 h day/10 h night photoperiod at 28.5 °C. Zebrafish larvae (2–4 dpf) and adult fish (3–12 months old) with similar sex ratio were analyzed for all experiments unless otherwise specified. Wild-type zebrafish strain used in this study was originally derived from Tubingen strain and has been maintained in our facility for more than 10 years by inbreeding. *Tg(fn1b:egfp*) (Shibata et al., 2018) was used in this study in addition to the Tgs established in this study.

Animal experimentation was performed in strict accordance with the recommendations in the Act on Welfare and Management of Animals in Japan and the Guide for the Care and Use of Laboratory Animals of the National Institutes of Health. All animals were handled according to the Animal Research Guidelines at Tokyo Institute of Technology. The protocol was approved by the Committee on the Ethics of Animal Experiments of the Tokyo Institute of Technology. All surgery was performed under 0.002% tricaine (3-aminobenzoic acid ethyl ester, Sigma-Aldrich, USA) anesthesia and every effort was made to minimize suffering.

### Analysis of transposable elements

The origin of the E1, E4, E5, and E6 enhancer sequences was identified based on the repeat annotation of the zebrafish genome assembly GRCz11/danRer11 available in the UCSC Genome Browser. The consensus sequences of the four TEs were retrieved from RepBase (Bao et al., 2015). The number of copies of the four TE families was determined using RepeatMasker with the rmblast search engine under the sensitive (-s) option (http://www.repeatmasker.org) using the zebrafish repeat library version 20181026 obtained from RepBase.

The numbers of copies in 1 Mbp windows of the zebrafish chromosomes were counted from the RepeatMasker output, and the TE densities were visualised with RIdeogram (Hao et al., 2020). For age distribution, the number of TE copies was counted for 1 % bin of TE sequence divergence from the consensus sequences based on the RepeatMasker output.

### Plasmid construction

For making a series of *fn1b* promoter constructs, the respective regions of *fn1b* were amplified by polymerase chain reaction (PCR) using KOD Plus Neo DNA polymerase (Toyobo) with primers containing XhoI and AgeI restriction enzyme (RE) sites on the respective primers. The amplified DNA was cloned into pCR Blunt II-TOPO (Thermo Fisher), excised with REs, and inserted into pT2KXIGDin (Urasaki et al., 2006). The *1*.*8k:egfp* construct was made from the *3*.*2k:egfp* by removing the sequence between Xho I and Pflm I. To facilitate the identification of transgenic fish, an *egfp* cassette under the control of the *crystalline alpha A* promoter (*cryaa:egfp*) was introduced at the Eco NI site, which lies upstream of the test DNA insertion site. The primers used for PCR are listed in Table S1.

E4, E5, and E2L were cloned from the *fn1b* BAC clone by PCR using primers containing RE sites. The *E4*-, *E5-*, and *E2L-miniP* constructs were created by inserting the cloned sequences in front of *miniP* and *egfp* (Shimizu et al., 2012). The *cryaa:egfp* cassette was introduced at the Eco NI site, upstream of the test DNA. To construct the plasmids with *fn1b* –0.7kb promoter, *miniP* was replaced with –0.7kb promoter at Sal I and Bam HI. The E2, E2S, E2S(dA), and E2S(dE) constructs were created from E2L by removing the partial sequences through PCR-mediated mutagenesis (KOD Plus Mutagenesis Kit, Toyobo).

To create constructs with the tandem repeat of E-box (*6xE*), AP-1 (*6xA*) or E-box/AP-1 (*6xE*-*6xA*), oligonucleotides were annealed and inserted in front of the *miniP* or –0.7kb promoters. HE1_DR1(m) was cloned from the zebrafish genome, and a randomly chosen clone was used to determine the sequence and create the constructs. All constructs were confirmed by sequencing.

### Zebrafish transgenesis and regeneration assay

Tol2 transgenesis (Kawakami, 2007) was used to generate the transgenic lines. Plasmid DNAs at a concentration of 25–40 ng/μL and 25 ng/μL of transposase mRNA were injected into fertilized eggs at one-cell stage.

Fin fold amputation of larvae was performed using a scalpel at 3–4 days post fertilization (dpf), as previously described (Kawakami et al., 2004). Fins of young (4–8 weeks post fertilization; hpf) or mature zebrafish (3–12 months post fertilization; mpf) were cut at approximately the middle of the fin. The regeneration response of the constructs that were positive in the F0 assay were further confirmed in more than two stable transgenic lines to avoid a possible positional effect of the integrated genomic site. To isolate F1 offspring with stable EGFP expression in response to fin amputation, F0 embryos were raised to adulthood and then randomly outcrossed or inbred to obtain the F1 carriers. In the case of *cryaa:egfp* cassette*-*containing constructs, the carriers were initially screened by the lens GFP fluorescence.

To induce scale regeneration, approximately 10 scales were removed from the trunk region of each fish using forceps. For the wound healing assay, the fin was punctured with a fine needle at the ray or inter-ray region.

### Heart resection

Heart resection was performed according to the procedure described before (Poss et al., 2002; Ito et al., 2014). Briefly, zebrafish of wild-type or *Tg(1*.*8k:egfp)* at 6-12 months old were anaesthetized in tricaine and placed ventral side up into a moist, slotted sponge. Using a micro scissor, a small incision was made in the area ventral to the position where the heart beat is visible. The chest of the fish was pressed with tweezers to completely expose the heart, and 20-30% of the lower part of the ventricle was removed with micro scissors. After surgery, fish were returned to water and stimulated to breathe by sending fresh water over the gills with a pipette. Fish were further recovered for 24 hrs in a tank with bubbling aeration. At 7 days after surgery, hearts were dissected and fixed with 4% (w/v) paraformaldehyde in PBS (PFA).

### Dominant negative lef1 (dnlef1) and heat shock experiment

The dominant negative form of the *lef1* plasmid, *pTol2(hsp70l:mKO2-2a-dnlef1)* (Akieda et al., 2019), was a generous gift from Toru Ishitani. A stable *Tg* line was generated at our facility. The line was established by screening the F1s for mKO2 fluorescence after a heat shock at 38 °C for 2 h. To examine the effect on fin regeneration, the first heat shock was performed 6 h before fin amputation.

### Search for TF-binding sites

The JASPER 2020 open access database (ver.7) (JASPER CORE Vertebra, https://jaspar.genereg.net/) (Fornes et al., 2020) was used for searching TF sites using default settings (relative score > 0.8). With some exception, only the potentially significant TF sites (highest Jasper score > 10.0) are shown in Fig. 3A–D and Fig. S3 A and 3B.

### *Xenopus* transgenic assay

The constructs, along with decondensed sperm nuclei and oocyte extracts, were injected into unfertilized eggs of *Xenopus laevis* (Kroll and Amaya, 1996). GFP fluorescence from the alpha-crystallin promoter was used to identify tadpoles with a higher transgenic efficiency. *Xenopus* stages were identified according to the Normal table of *Xenopus laevis* (Nieuwkoop and Faber, 1994). Amputation of the limb buds of tadpoles, under anaesthesia with tricaine, was performed at St. 52–54 using micro scissors, according to a previously published procedure (Hayashi et al., 2015).

### Whole-mount *in situ* hybridization

Whole-mount *in situ* hybridization (ISH) was performed according to the standard published protocol (Thisse and Thisse, 2008). A region of the EGFP coding sequence was amplified by PCR, and the product was used as a template to synthesize the RNA probe. The underlined part of the primer denotes the T7 promoter.

### Immunostaining and histological analysis

Zebrafish fins were fixed in PFA at 4 °C overnight, subsequently dehydrated with methanol, and stored at −20 °C. The stored samples were rehydrated with PBT (1× PBS, % Tween-20), equilibrated with 20 % (w/v) sucrose, embedded in OCT compound (Tissue-Tek, Sakura Finetek), and stored at −20 °C. Cryosections of 16–20 µm thickness were prepared for histological analysis.

For immunofluorescence staining, cryosections were washed twice with PBS and several times with PBT to remove residual OCT compound. Antibody staining was performed as previously described (Shibata et al., 2016). The GFP antibody (Nacalai Tesque or MBL) was used at 1:1000 dilution. The zns5 antibody was used at 1:100 dilution of the hybridoma supernatant (Developmental Studies Hybridoma Bank). The MF20 antibody (Invitrogen) was used at 1:1000 dilution. The sections were counterstained with 4′,6-diamidino-2-phenylindole (DAPI; 0.1 μg/mL, Invitrogen) and mounted with 80 % glycerol containing 25 mg/mL triethylenediamine (DABCO, Nacalai Tesque). Images were captured using a confocal microscope (Fluoview FV1000, Olympus).

### Chemical treatment

SP600125 (Selleck) was dissolved in dimethyl sulfoxide (DMSO) at 15 mM and stored at −80 °C. The stock solution was diluted to a working concentration with E3 buffer and administered to zebrafish at least 6 h before fin amputation. DMSO (0.1 %) was used as the vehicle control. The chemical solution was changed daily.

### Statistics

No statistical methods were used to determine sample size. Sample sizes were chosen based on previous publications and experiment types and are indicated in each Fig. legend. After selecting larvae or fish with wild type morphology, the clutch-mates were randomised into different groups for each experiment. No animal or sample was excluded from the analysis, unless the animal died during the procedure. Most assessments of phenotypes and expression patterns were replicated in at least three independent experiments except the heart regeneration (two replicates). Whenever possible, blinding was performed during data collection and analysis. In some experiments, when embryos had to undergo specific treatments, blinding was not possible, as the same investigator processed the samples and collected data. Sample sizes are indicated in the figures or legends.

For F0 assay of the regenerative-response, fin fold or fin of amputation was performed on animals with apparent EGFP expression in the lens, which reflects the rate of transgenesis. The numbers of larvae or fish with amputation-induced EGFP expression were scored; however, the ratios of EGFP response does not represent the strength of the enhancers, because the efficiency significantly varied in respective

injections. Throughout the Tg assays, all constructs that were judged as positive response in F0 assay were confirmed their response in more than two Tg lines. In case when it was difficult to judge the response in F0 assay, we repeated the injection and finally isolated the F1 carriers to confirm their responses.

## Acknowledgements

We are grateful to T. Ishitani for his generous gift of the plasmid, *pTol2(hsp70l:mKO2-2a-dnlef1)* and Kohei Ito for teaching us the detailed procedure of heart resection. We thank the Open Facility Centre for Life Science and Technology at the Tokyo Institute of Technology for sequencing and imaging support. E.S. was supported by a fellowship from the Education Academy of Computational Life Science at the Tokyo Institute of Technology. T.T. and T. Y. was supported by a Tsubame Scholarship for Doctoral Students at the Tokyo Institute of Technology.

## Author contributions

T. T. and T. Y. conceived the project, designed and performed most of the experiments, analyzed the data, and wrote the manuscript with input from all authors. E.S. designed and contributed to the initial parts of the study including transgenic assay. H.N. performed transposon analyses. H.O. performed experiments involving the *Xenopus* transgenic assay. A.K. supervised the study and wrote the manuscript with input from all authors.

## Data availability

All data reported in this paper will be shared by the lead contact upon request. This paper does not report original code. Any additional information required to reanalyze the data reported in this paper is available from the lead contact upon request.

## Supplementary information

Supplementary information available online at xxxxxxxxxxxx

## References

Akieda, Y., Ogamino, S., Furuie, H., Ishitani, S., Akiyoshi, R., Nogami, J., Masuda, T., Shimizu, N., Ohkawa, Y., and Ishitani, T. (2019). Cell competition corrects noisy Wnt morphogen gradients to achieve robust patterning in the zebrafish embryo. Nat. Commun. 10, 4710. doi: 10.1038/s41467-019-12609-4

Akimenko, M. A., Johnson, S. L., Westerfield, M., and Ekker, M. (1995). Differential induction of four msx homeobox genes during fin development and regeneration in zebrafish. Development 121, 347–357. doi: 10.1242/dev.121.2.347

Bao, W., Kojima, K. K., and Kohany, O. (2015). Repbase Update, a database of repetitive elements in eukaryotic genomes. Mobile DNA 6, 11. doi: 10.1186/s13100-015-0041-9

Bely, A. E., and Nyberg, K. G. (2010). Evolution of animal regeneration: re-emergence of a field. Trends Ecol. Evol. 25, 161–170. doi: 10.1016/j.tree.2009.08.005

Cadigan, K. M. and Waterman, M. L. (2012). TCF/LEFs and Wnt signaling in the nucleus. Cold Spring Harb. Perspect. Biol. 4, a007906. doi: 10.1101/cshperspect.a007906

Chuong, E. B., Elde, N. C., and Feschotte, C. (2016). Regulatory evolution of innate immunity through co-option of endogenous retroviruses. Science 351, 1083–1087. doi: 10.1126/science.aad5497

Fornes, O., Castro-Mondragon, J. A., Khan, A., van der Lee, R., Zhang, X., Richmond, P. A., Modi, B. P., Correard, S., Gheorghe, M., Baranašić, D. et al. (2020). JASPAR 2020: update of the open-access database of transcription factor binding profiles. Nucleic Acids Res. 48, D87–D92. doi: 10.1093/nar/gkz1001

Gauron, C., Rampon, C., Bouzaffour, M., Ipendey, E., Teillon, J., Volovitch, M., and Vriz, S. (2013). Sustained production of ROS triggers compensatory proliferation and is required for regeneration to proceed. Sci. Rep. 3, 2084. doi: 10.1038/srep02084

Goldman, J. A. and Poss, K. D. (2020). Gene regulatory programs of tissue regeneration. Nature Rev. Genet. 21, 511–525. doi:10.1038/s41576-020-0239-7

Hao, J., Ao, A., Zhou, L., Murphy, C. K., Frist, A. Y., Keel, J. J., Thorne, C. A., Kim, K., Lee, E., and Hong, C. C. (2013). Selective small molecule targeting β-catenin function discovered by in vivo chemical genetic screen. Cell Rep. 4, 898–904. doi: 10.1016/j.celrep.2013.07.047

Hao, Z., Lv, D., Ge, Y., Shi, J., Weijers, D., Yu, G., and Chen, J. (2020). RIdeogram: drawing SVG graphics to visualize and map genome-wide data on the idiograms. PeerJ Comput. Sci. 6, e251. doi: 10.7717/peerj-cs.251

Hayashi, S., Kawaguchi, A., Uchiyama, I., Kawasumi-Kita, A., Kobayashi, T., Nishide, H., Tsutsumi, R., Tsuru, K., Inoue, T., Ogino, H. et al. (2015). Epigenetic modification maintains intrinsic limb-cell identity in Xenopus limb bud regeneration. Dev. Biol. 406, 271–282. doi: 10.1016/j.ydbio.2015.08.013

Hermant, C., and Torres-Padilla, M.-E. (2021). TFs for TEs: the transcription factor repertoire of mammalian transposable elements. Genes Dev. 35, 22–39. doi: 10.1101/gad.344473.120

Hess, J., Angel, P., and Schorpp-Kistner, M. (2004). AP-1 subunits: quarrel and harmony among siblings. J. Cell Sci. 117, 5965–5973. doi: 10.1242/jcs.01589

Ishida, T., Nakajima, T., Kudo, A., and Kawakami, A. (2010). Phosphorylation of Junb family proteins by the Jun N-terminal kinase supports tissue regeneration in zebrafish. Dev. Biol. 340, 468–479. doi: 10.1016/j.ydbio.2010.01.036

Ito, K., Morioka, M., Kimura, S., Tasaki, M., Inohaya, K., and Kudo, A. (2014). Differential reparative phenotypes between zebrafish and medaka after cardiac injury. Dev. Dyn. 243, 1106–1115. doi: 10.1002/DVDY.24154

Ivanova, A. S., Tereshina, M. B., Ermakova, G. V., Belousov, V. V., and Zaraisky, A . G. (2013). Agr genes, missing in amniotes, are involved in the body appendages regeneration in frog tadpoles. Sci. Rep. 3, 1279. doi: 10.1038/srep01279

Jones, S. (2004). An overview of the basic helix-loop-helix proteins. Genome Biol. 5, 226. doi: 10.1186/gb-2004-5-6-226

Kang, J., Hu, J., Karra, R., Dickson, A. L., Tornini, V. A., Nachtrab, G., Gemberling, M., Goldman, J. A., Black, B. L., and Poss, K. D. (2016). Modulation of tissue repair by regeneration enhancer elements. Nature 532, 201–206. doi: 10.1038/nature17644

Kawakami, A., Fukazawa, T., and Takeda, H. (2004). Early fin primordia of zebrafish larvae regenerate by a similar growth control mechanism with adult regeneration. Dev. Dyn. 231, 693–699. doi: 10.1002/dvdy.20181

Kawakami, A. (2010). Stem cell system in tissue regeneration in fish. Dev. Growth Differ. 52, 77–87. doi: 10.1111/j.1440-169X.2009.01138.x

Kawakami, K. (2007). Tol2: a versatile gene transfer vector in vertebrates. Genome Biol. 8, S7. doi: 10.1186/gb-2007-8-s1-s7

Kroll, K. L., and Amaya, E. (1996). Transgenic Xenopus embryos from sperm nuclear transplantations reveal FGF signaling requirements during gastrulation. Development 122, 3173–3183. doi: 10.1242/dev.122.10.3173

Kumar, A., Godwin, J. W., Gates, P. B., Garza-Garcia, A. A., and Brockes, J. P. (2007). Molecular basis for the nerve dependence of limb regeneration in an adult vertebrate. Science 318, 772–777. doi: 10.1126/science.1147710

Lander, E. S., Linton, L. M., Birren, B., Nusbaum, C., Zody, M. C., Baldwin, J., Devon, K., Dewar, K., Doyle, M., FitzHugh, W. et al. (2001). Initial sequencing and analysis of the human genome. Nature 409, 860–921. doi: 10.1038/35057062

Lee, H. J., Hou, Y., Chen, Y., Dailey, Z. Z., Riddihough, A., Jang, H. S., Wang, T., and Johnson, S. L. (2020). Regenerating zebrafish fin epigenome is characterized by stable lineage-specific DNA methylation and dynamic chromatin accessibility. Genome Biology 21, 52. doi:10.1186/s13059-020-1948-0

Nieuwkoop, P. D., and Faber, J. (1994). Normal table of Xenopus laevis (Daudin): a systematical and chronological survey of the development from the fertilized egg till the end of metamorphosis. New York, U.S.A: Garland Science. doi:10.1201/9781003064565

Nishihara, H. (2019). Retrotransposons spread potential cis-regulatory elements during mammary gland evolution. Nucleic Acids Res. 47, 11551–11562. doi: 10.1093/nar/gkz1003

Nishihara, H. (2020). Transposable elements as genetic accelerators of evolution: contribution to genome size, gene regulatory network rewiring and morphological innovation. Genes Genet. Syst. 94, 269–281. doi: 10.1266/ggs.19-00029

Pfefferli, C. and Jaźwińska, A. (2017). The careg element reveals a common regulation of regeneration in the zebrafish myocardium and fin. Nat. Commun. 8, 15151. doi: 10.1038/ncomms15151

Phipps, L. S., Marshall, L., Dorey, K., and Amaya, E. (2020). Model Systems for Regeneration: Xenopus. Development 147, dev180844. doi: 10.1242/dev.180844

Poss, D., Wilson, L. G., and Keating, M. T. (2002). Heart Regeneration in Zebrafish. Science 298, 2188–2190. doi: 10.1126/science.1077857

Poss, K. D. (2010). Advances in understanding tissue regenerative capacity and mechanisms in animals. Nat. Rev. Genet. 11, 710–722. doi: 10.1038/nrg2879

Shibata, E., Yokota, Y., Horita, N., Kudo, A., Abe, G., Kawakami, K., and Kawakami, A. (2016). Fgf signalling controls diverse aspects of fin regeneration. Development 143, 2920–2929. doi: 10.1242/dev.140699

Shibata, E., Ando, K., Murase, E., and Kawakami, A. (2018). Heterogeneous fates and dynamic rearrangement of regenerative epidermis-derived cells during zebrafish fin regeneration. Development 145, dev162016. doi: 10.1242/dev.162016

Shimizu, N., Kawakami, K., and Ishitani, T. (2012). Visualization and exploration of Tcf/Lef function using a highly responsive Wnt/β-catenin signaling-reporter transgenic zebrafish. Dev. Biol. 370, 71–85. doi: 10.1016/j.ydbio.2012.07.016

Tanaka, E. M. (2016). The Molecular and Cellular Choreography of Appendage Regeneration. Cell 165, 1598–1608. doi: 10.1016/j.cell.2016.05.038

Thisse, C. and Thisse, B. (2008). High-resolution in situ hybridization to whole-mount zebrafish embryos. Nat. Protocol 3, 59–69. doi: 10.1038/nprot.2007.514

Thompson, J. D., Ou, J., Lee, N., Shin, K., Cigliola, V., Song, L., Crawford, J. E., Kang, J., and Poss, K. D. (2020). Identification and requirements of enhancers that direct gene expression during zebrafish fin regeneration. Development 47, dev191262. doi:10.1242/dev.191262

Urasaki, A., Morvan, G., and Kawakami, K. (2006). Functional dissection of the Tol2 transposable element identified the minimal cis-sequence and a highly repetitive sequence in the subterminal region essential for transposition. Genetics 174, 639–649. doi: 10.1534/genetics.106.060244

Vacik, T. and Lemke, G. (2011). Dominant-negative isoforms of Tcf/Lef proteins in development and disease. Cell Cycle 10, 4199–4200. doi: 10.4161/cc.10.24.18465

Visel, A., Rubin, E. M., and Pennacchio, L. A. (2009). Genomic views of distant-acting enhancers. Nature 461, 199–205. doi: 10.1038/nature08451

Wang, J, Karra, R., Dickson, A. L., and Poss, K. D. (2013). Fibronectin is deposited by injury-activated epicardial cells and is necessary for zebrafish heart regeneration. Dev. Biol. 382, 427–435. doi: org/10.1016/j.ydbio.2013.08.012

Wang, L. H., and Baker, N. E. (2015). E proteins and ID proteins: Helix-Loop-Helix Partners in Development and Disease. Dev. Cell 35, 269–280. doi: 10.1016/j.devcel.2015.10.019

Wang, W., Hu, C. K., Zeng, A., Alegre, D., Hu, D., Gotting, K., Ortega-Granillo, A., Wang, Y., Robb, S., Schnittker, R. et al. (2020). Changes in regeneration-responsive enhancers shape regenerative capacities in vertebrates. Science 369, eaaz3090. doi: 10.1126/science.aaz3090

Yoshinari, N., Ishida, T., Kudo, A., and Kawakami, A. (2009). Gene expression and functional analysis of zebrafish larval fin fold regeneration. Dev. Biol. 325, 71–81. doi: 10.1016/j.ydbio.2008.09.028

